# An FEV-Associated Metastasis-Initiating-Like State Linked to Tumor Cell Plasticity and Immune Niche Remodeling in Prostate Cancer

**DOI:** 10.64898/2026.09.23.753952

**Authors:** Sijia Wu, Jiangpeng Wei, Xiaorui Shi, Xin Liu, Xiaobo Zhou, Liyu Huang

## Abstract

Prostate cancer progression is accompanied by dynamic remodeling of malignant cell states, whose metastatic potential and regulatory programs remain poorly defined. Here, we integrated single-cell, spatial, bulk transcriptomic, epigenomic, and experimental data to characterize malignant cell state transitions and identify candidate regulators associated with metastatic progression. A distinct intermediate state exhibits increased copy-number alterations, stemness, proliferation, and MYC activity, together with the highest transcriptional similarity to metastatic tumor cells, and is therefore designated as a metastasis-initiating-like (MIC-like) state. Transcriptional and regulon dynamics along the malignant cell state trajectory highlight FEV as a prominent candidate associated with the MIC-like state, with concordant expression and regulon activity patterns. Functionally, FEV is associated with a hybrid epithelial-mesenchymal phenotype, and its perturbation disrupts this phenotype, accompanied by reduced epithelial and mesenchymal features as well as impaired prostate cancer cell migration and invasion. Beyond this tumor-intrinsic feature, FEV is also associated with reduced cytotoxicity and interferon responses in the tumor microenvironment through its regulation of MIF-family genes. In addition to its potential roles in tumor cell plasticity and immune remodeling, FEV shows a reciprocal association with androgen signaling and distinct expression patterns across treatment response groups, highlighting its potential relevance in therapeutic contexts.

## 1. Introduction

Prostate cancer is one of the most frequently diagnosed malignancies in men and a major cause of cancer-related mortality worldwide [1]. Although many patients with localized disease can be effectively managed, a subset of tumors progresses to locally advanced, metastatic, and ultimately castration-resistant disease, which remains a major cause of prostate cancer–related mortality. This progression involves the acquisition of malignant features that enable tumor cells to survive, adapt, and disseminate beyond the primary site [2]. Understanding how tumor cells evolve during prostate cancer progression is therefore essential for elucidating the mechanisms underlying disease advancement and metastasis.

The emergence of single-cell and spatial transcriptomic technologies has substantially expanded our understanding of cellular heterogeneity in prostate cancer [3-5]. In primary and advanced tumors, accumulating evidence has revealed distinct malignant cell states characterized by differences in androgen receptor (AR) activity [6], mesenchymal and stem-like features [7], and interactions with the immune microenvironment [8]. These findings suggest that prostate cancer progression is accompanied by dynamic remodeling of tumor cell states, with distinct cellular states potentially representing different stages along the progression toward metastatic disease [9]. However, how these distinct malignant states are organized along the trajectory of disease progression remains incompletely understood. In particular, identifying tumor cell states that emerge before the acquisition of overt metastatic phenotypes may provide important insights into the cellular processes underlying metastatic progression.

Transcription factors play central roles in establishing and maintaining tumor cell states by coordinating gene expression programs involved in proliferation, differentiation, lineage plasticity, and tumor progression. In prostate cancer, androgen receptor (AR) functions as a central lineage-defining transcription factor and a major driver of tumor-cell growth and survival [6]. Its transcriptional activity is shaped by a network of lineage-defining and pioneer factors, including FOXA1, HOXB13, and GATA2. FOXA1 facilitates chromatin accessibility and helps establish AR binding landscapes [10], whereas HOXB13 [11] and GATA2 [12] contribute to the regulation and reprogramming of AR-associated transcriptional programs. Together, these factors illustrate how transcriptional and epigenetic regulatory networks can shape prostate cancer cell identity and disease progression. Emerging evidence has also implicated FEV, an ETS-family transcription factor, in prostate cancer progression. FEV expression has been reported to be reduced in recurrent and metastatic prostate cancer, with functional studies linking its reduced expression to tumor-suppressive effects on prostate cancer cell proliferation, migration, invasion, and tumor growth [13]. Nevertheless, whether FEV and other transcriptional regulators define specific tumor cell states along the trajectory of prostate cancer progression, and how these regulatory programs relate to metastatic potential, remain incompletely defined. In this study, we integrated single-cell, spatial, bulk transcriptomic, epigenomic, and experimental datasets to characterize tumor cell heterogeneity and identify regulatory programs associated with a metastatic intermediate state in prostate cancer.

## 2. Results

### 2.1 Identification of a metastasis-initiating-like tumor subtype in prostate cancer

Tumor cells exhibit substantial heterogeneity, which may underlie distinct functional states and clinical behaviors. To characterize tumor cell heterogeneity in prostate cancer, we analyzed a single-cell transcriptomic dataset and identified 863 malignant epithelial cells based on epithelial marker expression and inferred copy-number variation (CNV) patterns (Figure S1). Unsupervised clustering resolved eight distinct tumor subtypes (Figure 1A), which occupied distinct positions along a pseudotime trajectory (Figure 1B), suggesting a continuum of tumor cell states.

**Figure 1.**
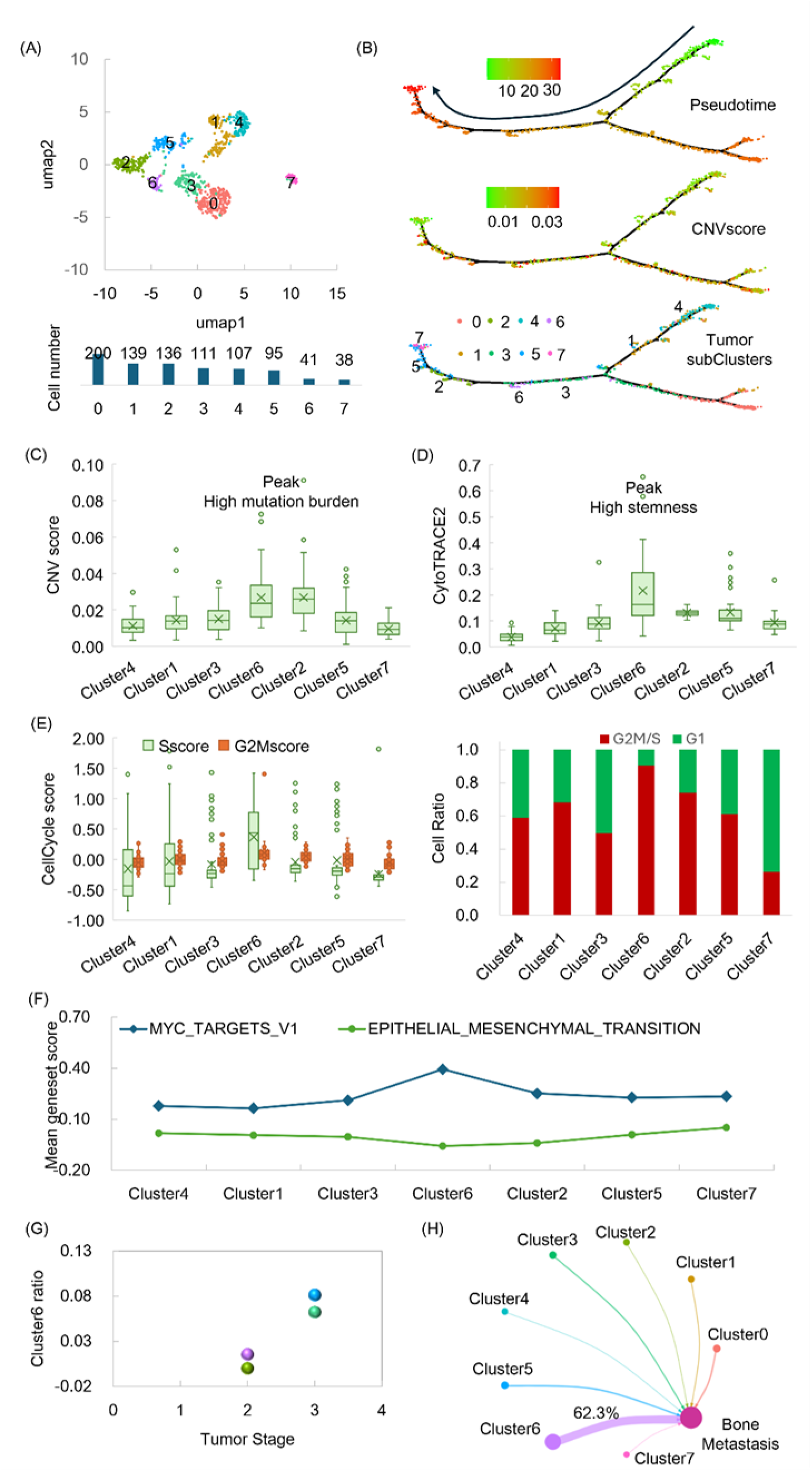
Identification of a metastasis-initiating-like tumor subtype in prostate cancer. (A) UMAP visualization of tumor cell subtypes, with the number of cells in each subtype shown below. (B) Pseudotime trajectory of the tumor cells inferred using Monocle. (C) Distribution of copy number variation (CNV) scores across tumor subtypes along the pseudotime trajectory estimated using inferCNV. (D) Distribution of CytoTRACE2 scores across tumor subtypes along the pseudotime trajectory. (E) S-phase and G2/M-phase scores and the proportion of cells in the G1, S, and G2/M phases across tumor subtypes, calculated using CellCycleScoring. (F) Enrichment scores for the MYC target and epithelial-mesenchymal transition hallmark gene sets across tumor subtypes, calculated using AddModuleScore. (G) Proportion of cluster 6 cells in four primary prostate tumors across different tumor stages. (H) Proportion of metastatic tumor cells showing the highest transcriptomic similarity in principal component analysis (PCA) space to each primary tumor subtype, determined using K-nearest neighbors (KNN).

Cluster 6 was positioned near the central region of the trajectory and exhibited a distinct malignant phenotype. Compared with the other tumor subtypes, cluster 6 shows the highest CNV score (Figure 1C) indicating increased genomic instability, and the highest stemness score (Figure 1D). Cluster 6 also displays the strongest proliferative phenotype, as indicated by both cell-cycle scores, the proportion of cycling cells (Figure 1E) and the scores of MYC TARGETS pathway (Figure 1F). Together, these features suggest that cluster 6 represents a highly malignant, stem-like, and proliferative tumor cell state.

The pseudotime trajectory further suggests a progressive transition toward this malignant state. Tumor cells locate between the root and cluster 6 shows a gradual increase in malignant features, culminating in the highly proliferative and stem-like phenotype observed in cluster 6. In contrast, tumor cells extending from cluster 6 toward the terminal state exhibit progressively higher epithelial–mesenchymal transition (EMT) scores (Figure 1F), suggesting that the cluster 6 state may represent an intermediate state preceding acquisition of a more mesenchymal phenotype.

The clinical and metastatic distributions of cluster 6 further support its potential relevance to prostate cancer progression. Among the four primary prostate cancer samples, tumors with higher pathologic stage tend to contain a greater proportion of cluster 6 cells (Figure 1G). Moreover, when tumor cells from primary and metastatic lesions were compared based on transcriptomic similarity, more than 60% of metastatic tumor cells show their highest similarity to cells within cluster 6 (Figure 1H). Collectively, these findings identify cluster 6 as a metastasis-initiating-like (MIC-like) tumor cell state in prostate cancer, which is subsequently investigated to define its molecular determinants and potential clinical relevance.

### 2.2 Transcriptional reprogramming of the metastasis-initiating-like tumor subtype

To further characterize the molecular features of cluster 6 tumor cells, we identified 1,357 differentially expressed genes between cluster 6 and the other tumor subtypes (Figure 2A). Functional enrichment analysis reveals a distinct transcriptional program in cluster 6 (Figure 2B-C). Genes upregulated in cluster 6 are predominantly enriched in protein synthesis, mitochondrial metabolism, DNA replication, and cell-cycle progression, whereas downregulated genes are mainly associated with cell adhesion and immune-related processes. Together, these findings indicate that tumor cells in cluster 6 acquire a transcriptional state characterized by enhanced biosynthetic, metabolic, and proliferative activity, accompanied by attenuated immune-interacting programs.

**Figure 2.**
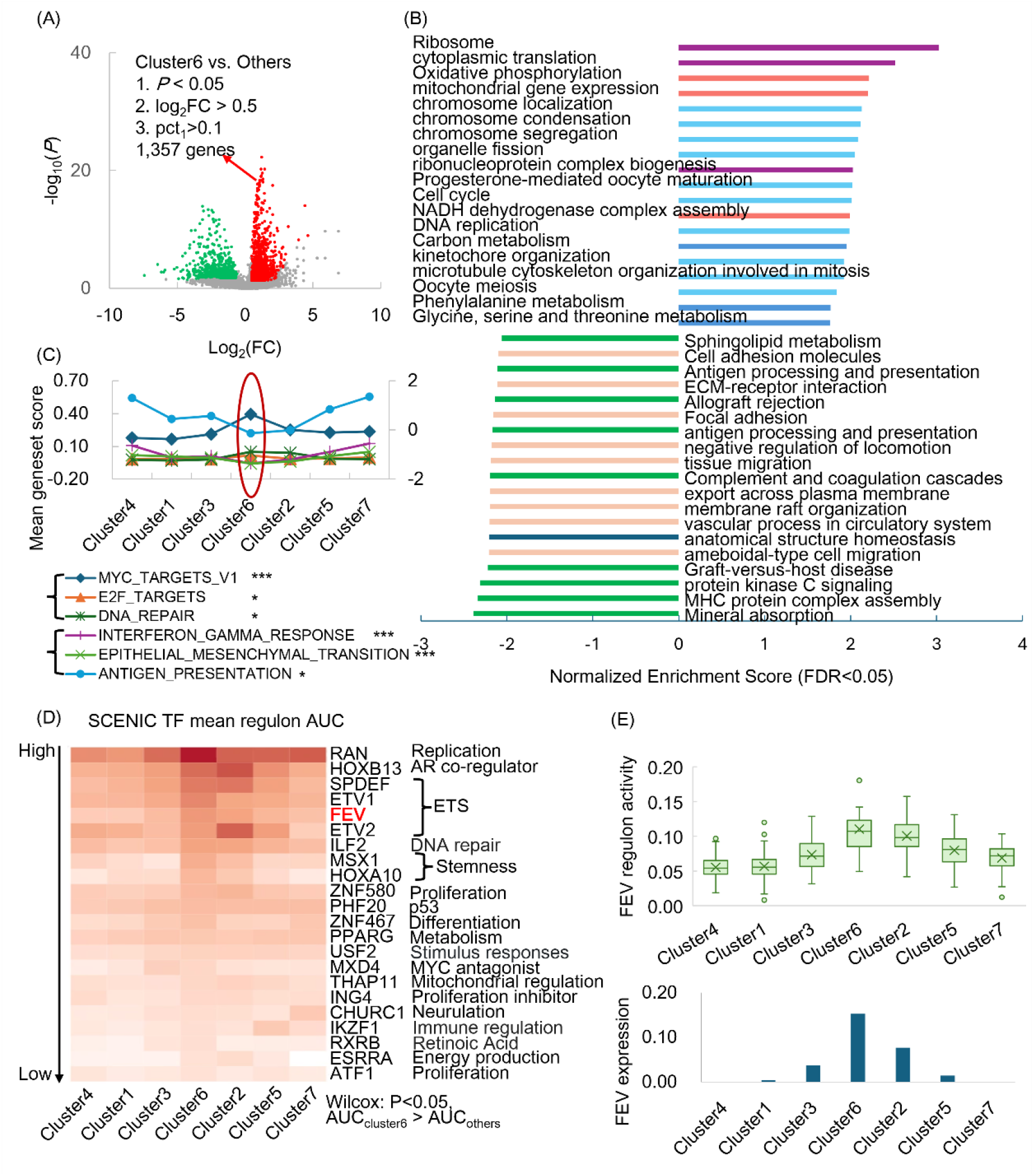
Transcriptional reprogramming of the metastasis-initiating-like tumor subtype. (A) Differentially expressed genes between cluster 6 and the other tumor subtypes identified using FindMarkers. (B) Gene set enrichment analysis (GSEA) of differentially expressed genes associated with cluster 6 performed using WebGestalt. (C) Representative pathway activities calculated using AddModuleScore across tumor subtypes ordered along the pseudotime trajectory, with activity levels in cluster 6 compared with those in the other subtypes using the Wilcoxon rank-sum test. (D) Transcription factor (TF) regulons with significantly higher activity in cluster 6 than in the other tumor subtypes, ranked by their mean regulon activity in cluster 6. (E) FEV expression and regulon activity across tumor subtypes ordered along the pseudotime trajectory. AUC: area under the curve; ***: *P* < 0.001; **: *P* < 0.01; *: *P* < 0.05.

The genes associated with the acquisition and subsequent transition of the cluster 6 state were further examined that whose expression increased from the trajectory root toward cluster 6 and subsequently decreased toward the terminal state. Among the significantly altered genes in cluster 6, 80 genes exhibit this characteristic trajectory-dependent expression pattern (Figure S2). These genes encompass multiple functional categories, including transcriptional regulation, hypoxia and cellular stress responses, androgen receptor (AR)-related signaling, mitochondrial energy metabolism, metabolic reprogramming, cell-cycle regulation, differentiation and proliferation, signal transduction and intracellular trafficking, and protein synthesis and proteostasis. Thus, these genes collectively define a broad transcriptional program associated with the emergence of the MIC-like tumor state.

Further, transcription factor (TF) activities were investigated to identify potential regulators underlying the transcriptional reprogramming associated with the MIC-like state. A total of 22 TFs shows significantly higher regulon activity in cluster 6 than in the other tumor subtypes (Figure 2D). Notably, only two TFs, FEV and MXD4, overlap with the 80 trajectory-associated genes identified above (Figure S2). FEV was prioritized for further investigation because both its expression and regulon activity closely followed the trajectory of the MIC-like state, progressively increasing from the trajectory root toward cluster 6 and subsequently decreasing toward the terminal state (Figure 2E). Importantly, the increased expression of FEV along the malignant trajectory was independently validated in an additional single-cell transcriptomic dataset (Figure S3A), three bulk transcriptomic datasets (Figure S3B), and a spatial transcriptomic dataset (Figure S3C). Collectively, these findings identify FEV as a candidate transcriptional regulator associated with the MIC-like tumor state, supporting further investigation of its biological and clinical relevance in prostate cancer.

### 2.3 FEV is associated with a hybrid epithelial–mesenchymal state

Metastatic dissemination is thought to be initiated by metastasis-initiating cancer cells with stem-like and immune-evasive properties [14], features that characterize the cluster 6 tumor state identified above. Intravasation represents an early step in metastatic dissemination and requires tumor cells to acquire increased migratory and invasive capacities, processes closely associated with EMT [15]. Given the high expression of FEV in cluster 6, the relationship between FEV expression and EMT was subsequently investigated. Along the trajectory from cluster 6 toward the terminal cluster 7, the VIM/CDH1 ratio progressively increases, consistent with a transition toward a more mesenchymal state, whereas FEV expression shows a reciprocal decrease (Figure 3A). Notably, the relatively high FEV expression in cluster 6, together with its intermediate epithelial-mesenchymal phenotype, suggests that this MIC-like state may represent a hybrid epithelial – mesenchymal state rather than a fully epithelial or mesenchymal phenotype. This pattern is also observed in the cancer cell line encyclopedia (CCLE) dataset, where 22Rv1 cells exhibits an intermediate CDH2/CDH1 ratio and relatively high FEV expression, whereas cell lines with progressively higher CDH2/CDH1 ratios generally shows lower FEV expression (Figure 3B). Thus, both the single-cell trajectory and cell-line data suggest that FEV expression is preferentially maintained in a hybrid epithelial–mesenchymal state and decreases as cells acquire a more mesenchymal phenotype.

**Figure 3.**
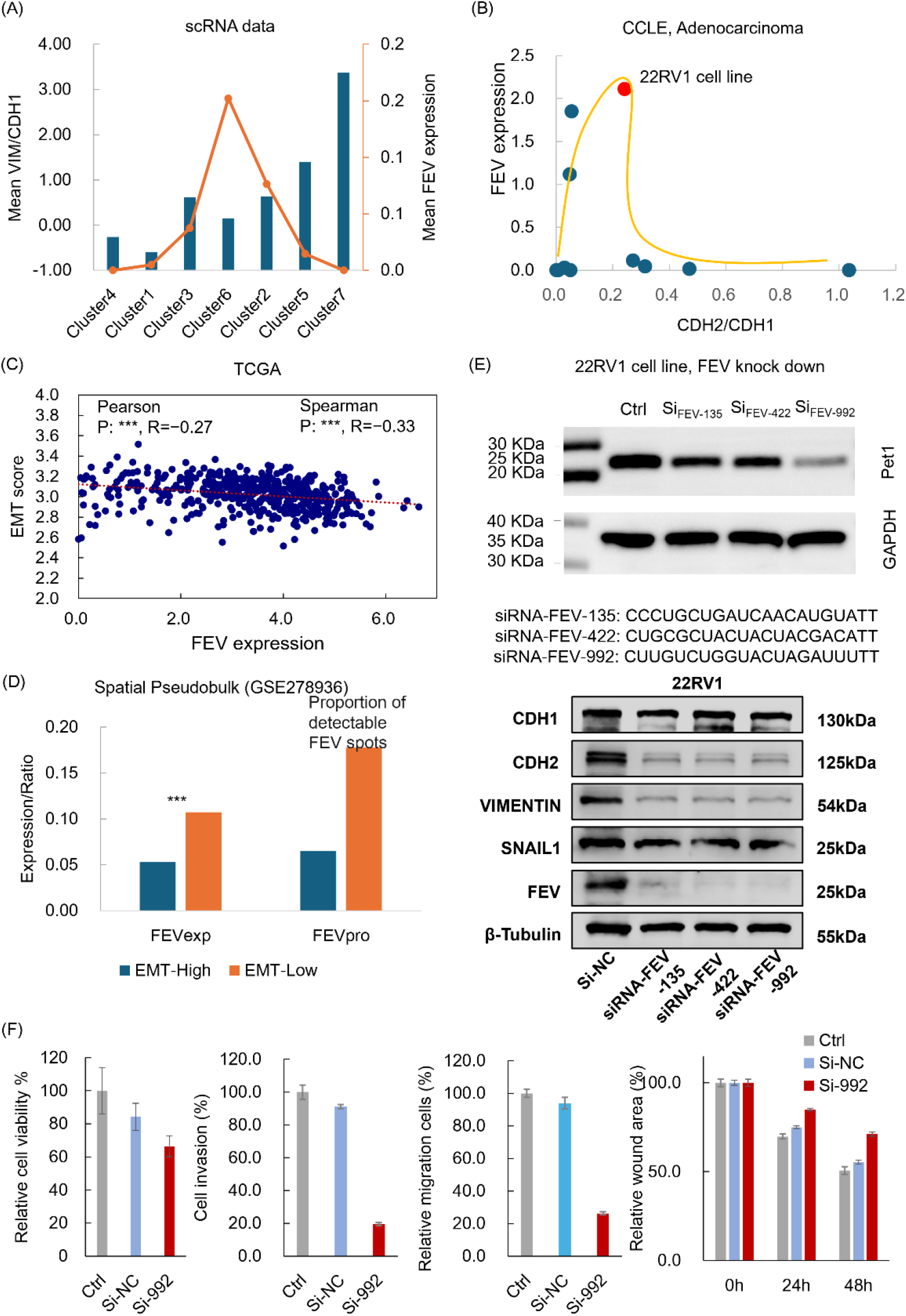
Association between FEV expression and epithelial–mesenchymal transition (EMT). (A) Mean VIM/CDH1 ratios and FEV expression across tumor subtypes ordered along the pseudotime trajectory. (B) CDH2/CDH1 ratios and FEV expression across prostate cancer cell lines in the cancer cell line encyclopedia (CCLE)database. (C) Association between FEV expression and EMT gene set scores in The Cancer Genome Atlas (TCGA) dataset, assessed using GSVA. (D) Comparison of FEV expression and the proportion of spots with detectable FEV expression between spots with high and low EMT scores in a spatial transcriptomic dataset, using the Wilcoxon rank-sum test. (E) Expression of epithelial and mesenchymal markers in 22Rv1 cells following FEV knockdown. (F) Comparison of the cell viability, invasion, and migratory capacities of 22Rv1 cells following FEV knockdown, assessed using cell viability, transwell invasion, and wound-healing assays.

This relationship is further supported by independent bulk transcriptomic datasets, in which FEV expression is significantly negatively correlated with EMT enrichment scores (Figure 3C and Figure S3D). Spatial transcriptomic analyses provide an additional layer of validation, showing that both FEV expression and the proportion of spots with detectable FEV expression are reduced in regions with higher EMT scores across three independent datasets (Figure 3D and Figure S3E). Collectively, these cross-platform analyses suggest that FEV is preferentially expressed in a hybrid epithelial–mesenchymal state associated with the MIC-like phenotype and declines during progression toward a more mesenchymal state.

To further examine whether FEV contributes to the maintenance of this phenotype, FEV was knocked down in 22Rv1 cells. FEV depletion resulted in coordinated reductions in both epithelial and mesenchymal markers (Figure 3E). It suggests that FEV loss does not simply promote or inhibit a conventional epithelial-to-mesenchymal transition but instead disrupts the transcriptional program underlying the hybrid epithelial–mesenchymal state. Consistent with this interpretation, FEV knockdown significantly reduced cell motility and invasive capacity in both scratch-wound and Transwell assays (Figure 3F and Figure S4). These functional effects are consistent with the transcriptomic observations across single-cell, bulk, cell-line, and spatial datasets, collectively supporting a role for FEV in maintaining the plastic malignant phenotype and functional properties of the MIC-like tumor state. Thus, FEV loss compromises the malignant phenotype of 22Rv1 cells and attenuates their migratory and invasive capacities.

### 2.4 FEV regulates MIF family genes in prostate cancer

Given the potential role of FEV in prostate cancer progression, further investigation was conducted to characterize its downstream regulatory functions. SCENIC analysis identifies a broad spectrum of putative FEV target genes involved in cell cycle regulation, intracellular transport, ubiquitination, RNA processing, translation, and transcriptional regulation (Figure S5A), consistent with potential roles in tumor cell proliferation. In addition, FEV targets were enriched in biological processes related to redox homeostasis, DNA damage and repair, and metabolism (Figure S5A), which may contribute to tumor microenvironment interactions. Among the predicted targets, PCGEM1, a well-characterized prostate cancer-specific lncRNA implicates in early prostate carcinogenesis, proliferation, invasion, and migration, is also identified as a potential FEV target [16].

Considering the critical role of immune evasion in MIC-like cells, the potential involvement of FEV in tumor–immune interactions was further examined. Among the predicted FEV targets, DDT (D-dopachrome tautomerase) belongs to the macrophage migration inhibitory factor (MIF) family, which plays important roles in immune regulation and tumor microenvironment interactions [17]. The relationship between FEV and the MIF family genes MIF and DDT was therefore examined across multiple transcriptomic datasets. Higher FEV expression is consistently associated with increased DDT and MIF expression across a single-cell transcriptomic dataset, two independent bulk transcriptomic datasets, a prostate cancer cell line dataset, and a spatial transcriptomic dataset (Figure 4A).

**Figure 4.**
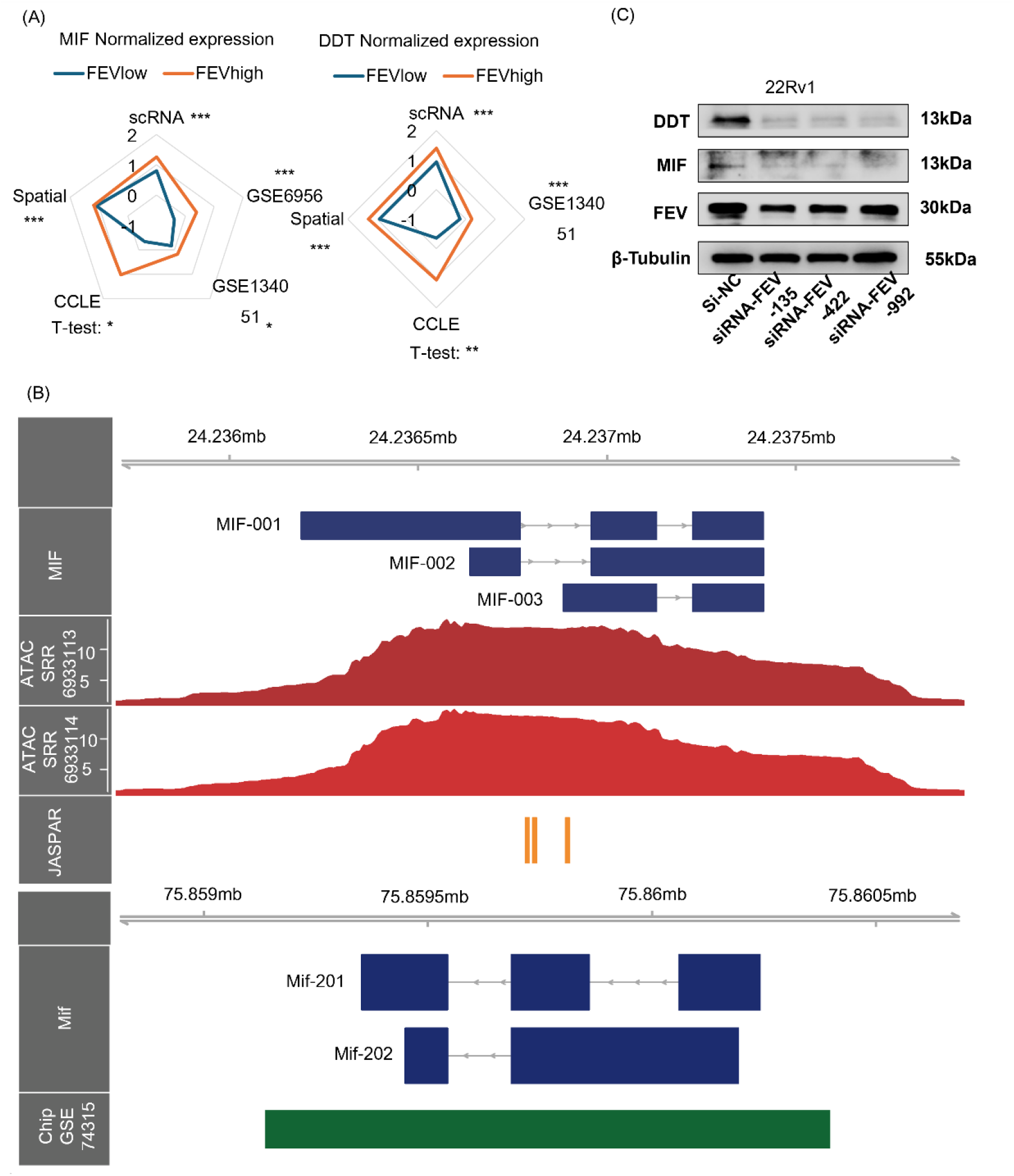
FEV regulates MIF family genes in prostate cancer. (A) Comparison of MIF and DDT expression between FEV-high and FEV-low groups across multiple multi-omics datasets, using the Wilcoxon rank-sum test for unannotated comparisons and the Student’ s t-test for comparisons indicated in the figure. (B) Accessible chromatin regions identified by ATAC-seq, putative FEV-binding sites predicted by JASPAR motif analysis, and FEV-binding signals identified by ChIP-seq analysis at the MIF locus. (C) Expression of MIF and DDT following FEV knockdown in 22Rv1 cells.

The potential regulatory relationship between FEV and MIF family genes was further evaluated using epigenomic analyses. In 22Rv1 cells, ATAC-seq reveals accessible chromatin regions surrounding the MIF and DDT loci (Figure 4B and Figure S5B). Motif analysis using JASPAR [18] identifies putative FEV binding motifs within these accessible chromatin regions, suggesting that FEV may directly regulate MIF and DDT transcription. Consistently, publicly available ChIP-seq data for mouse Pet-1, the ortholog of human FEV, reveals prominent binding peaks overlapping the MIF locus (Figure 4B). Collectively, these multi-omics findings support a potential regulatory role of FEV in controlling MIF family genes, particularly MIF, and suggest a possible mechanism through which FEV may influence tumor immune interactions in prostate cancer.

To further validate the regulatory association between FEV and MIF family genes, FEV was knocked down in 22Rv1 cells, followed by assessment of MIF and DDT expression. FEV knockdown resulted in a significant reduction in the expression of both MIF and DDT (Figure 4C). Together, these findings provide complementary evidence supporting a regulatory association between FEV and MIF family genes in prostate cancer.

### 2.5 FEV reshapes the immune microenvironment through regulation of MIF

Given that FEV regulates MIF expression in epithelial tumor cells, its potential role in shaping the immune microenvironment was further investigated. Cell–cell communication analysis was performed across tumor subclusters along the trajectory. Notably, tumor subclusters locate in the middle of the trajectory exhibits relatively limited signaling interactions with the surrounding microenvironment (Figure 5A), potentially reflecting a state of enhanced immune evasion. Among the identified interactions, the MIC-like cell subtype shows prominent MIF-mediated signaling through receptor complexes comprising CD74 and CXCR4 or CD44 (Figure 5B). These strong interactions are consistent with the highest MIF expression and MIF signaling interaction score observed in the MIC-like cell subtype (Figure 5C–D). MIF-mediated signaling is particularly prominent between the MIC-like cell subtype and immune cells (Figure 5E). At the patient level, the abundance of the MIC-like cell subtype is negatively associated with immune cytotoxicity and Interferon response (Figure 5F). The results present an altered immune microenvironment associated with FEV and the MIC-like status.

**Figure 5.**
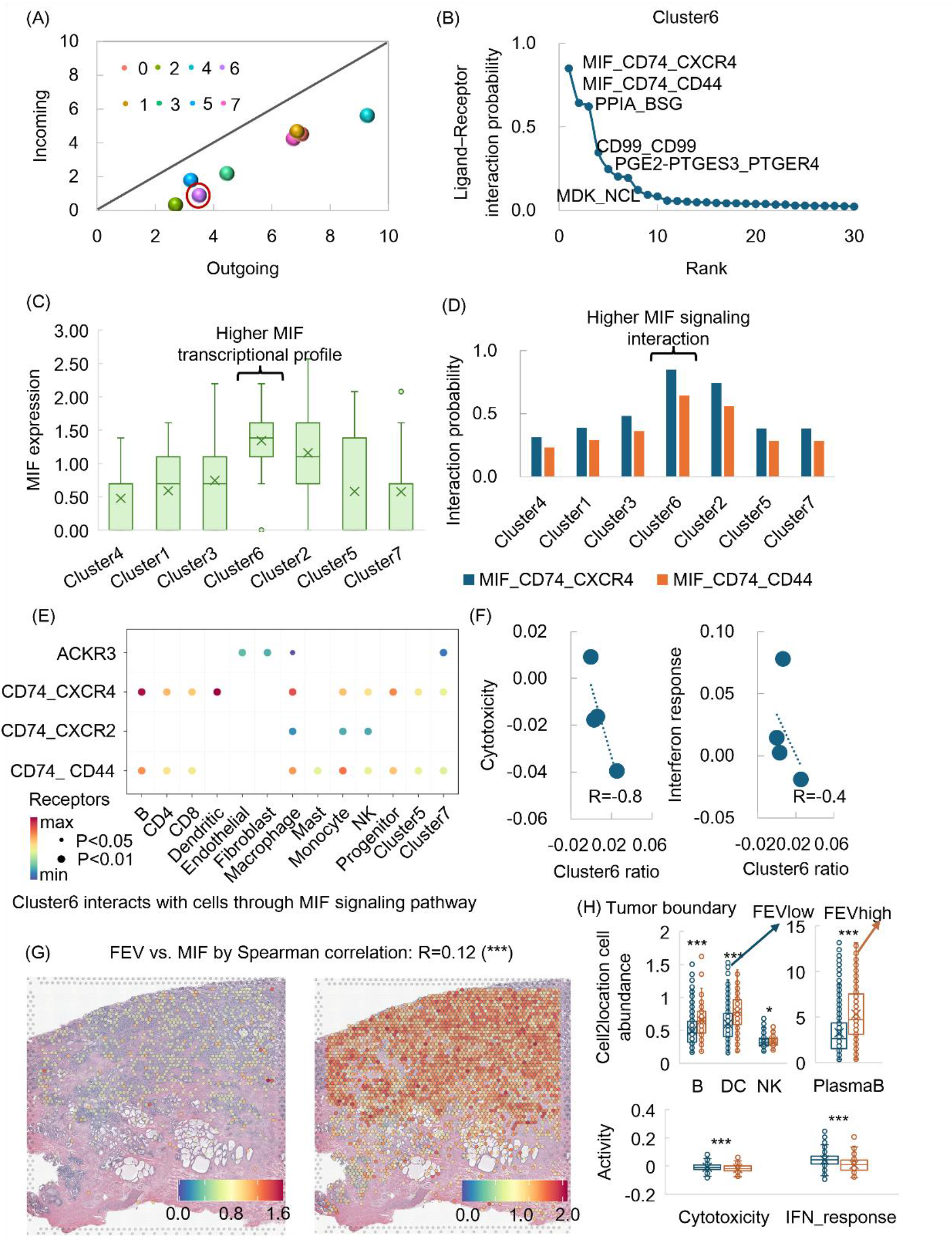
Metastasis-initiating-like cells with higher FEV expression are associated with immune niche remodeling through MIF-associated signaling. (A) Outgoing and incoming signaling strength of different tumor subclusters estimated using CellChat. (B) Ranked ligand–receptor interactions in tumor subcluster 6. (C) MIF expression across tumor subclusters along the pseudotime trajectory. (D) Summed MIF signaling interaction probabilities across tumor subclusters along the pseudotime trajectory. (E) MIF signaling interaction probability between MIC-like cells and other cell types. (F) Associations of cytotoxicity and interferon response score, calculated using AddModuleScore, with the proportion of cluster 6 cells at the patient level. (G) Associations between FEV and MIF expression within individual spatial transcriptomic slide. (H) Comparison of immune cell abundances estimated using cell2location and immune activity scores between FEV-high and FEV-low spots at the tumor boundary.

Consistent with the associations between FEV and MIF observed in spatial transcriptomic data (Figure 4A, Figure 5G, and Figure S6A), their spatial relationship with the immune microenvironment was further examined to characterize the immune niche associated with the FEV- and MIC-like states. FEV-enriched spots are located farther from the tumor boundary (Figure S6B–C) and preferentially localized within the tumor core, a region implicated in the localization of metastatic seeds [19]. Focusing on tumor-boundary spots, FEV-high spots are surrounded by higher abundances of immune cell populations, yet exhibit reduced cytotoxicity and interferon response scores compared with FEV-low spots (Figure 5H and Figure S6D). Together, these findings support a spatially remodeled immune niche associated with the FEV-high and MIC-like tumor states.

### 2.6 FEV expression is associated with AR signaling during prostate cancer progression

Androgen receptor (AR) signaling plays a central role in prostate cancer progression and therapeutic response. The relationship between AR activity and FEV expression was therefore examined across prostate cancer models and clinical specimens. In single-cell transcriptomic data, changes in FEV expression are closely accompanied by alterations in AR activity across tumor cell states (Figure 6A). Consistently, three independent bulk primary prostate cancer transcriptomic cohorts demonstrate significant positive associations between AR activity and FEV expression (Figure 6B and Figure S7A). This association is further supported by analyses of primary prostate cancer cell lines, in which lower AR activity is generally accompanied by reduced FEV expression (Figure 6C).

**Figure 6.**
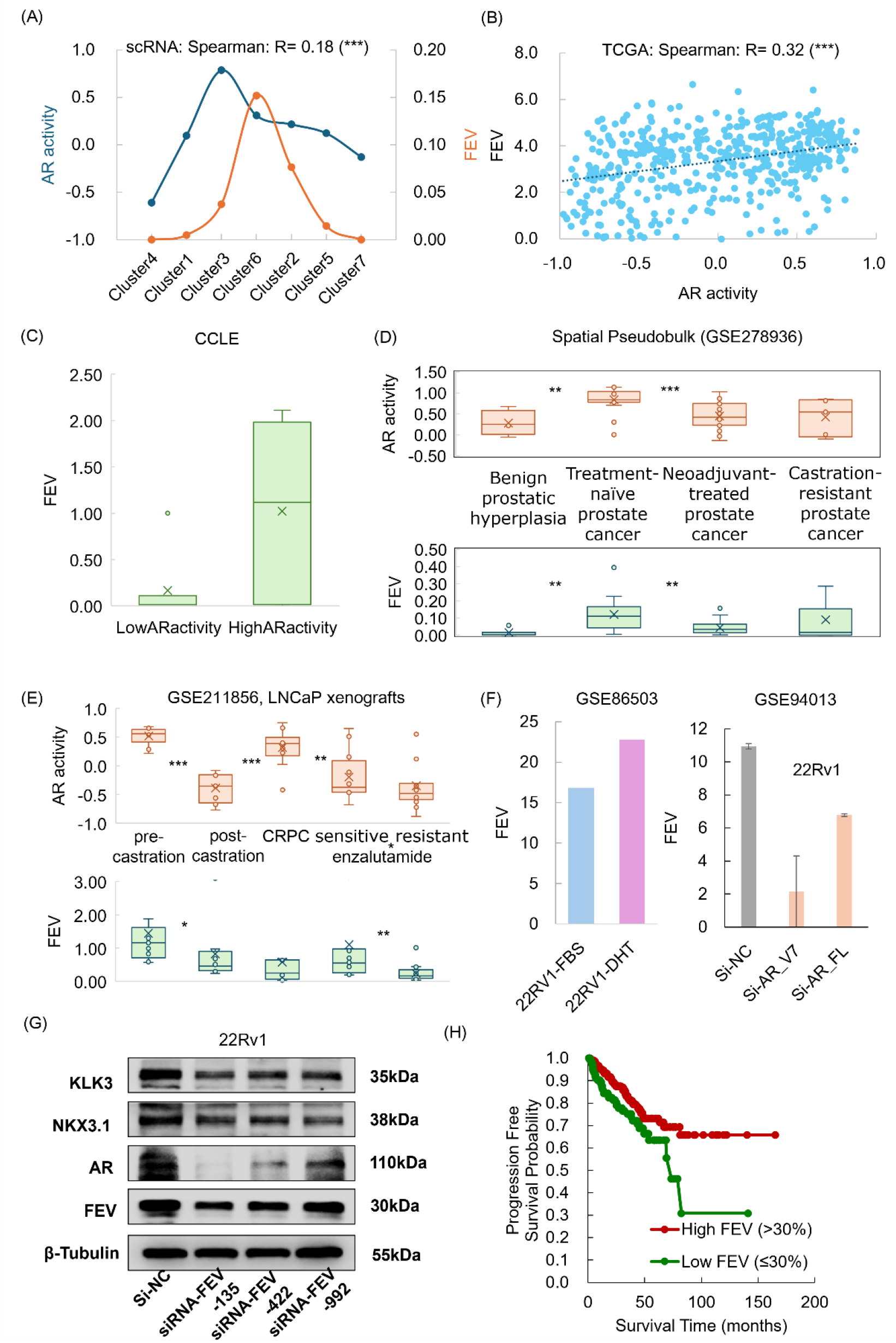
Associations between AR activity and FEV expression. (A) AR activity and FEV expression along the trajectory of tumor cells in the scRNA-seq dataset. (B) Association between AR activity and FEV expression in the TCGA prostate cancer dataset. (C) Comparison of FEV expression between prostate cancer cell lines with low and high AR activity. (D) AR activity and FEV expression across different stages of primary prostate cancer progression in spatial transcriptomic datasets. (E) AR activity and FEV expression in LNCaP xenografts across different stages of primary prostate cancer progression. (F) FEV expression in 22Rv1 cells following DHT treatment or knockdown of AR-FL or AR-V7. (G) The expressions of AR and AR target genes in 22Rv1 cells following FEV knockdown.(H) Survival analysis according to FEV expression in the TCGA prostate cancer dataset

The relationship between AR activity and FEV expression was further examined across different stages of prostate cancer progression. In spatial transcriptomic datasets encompassing benign prostatic hyperplasia, treatment-naïve prostate cancer, neoadjuvant-treated prostate cancer, and castration-resistant prostate cancer (CRPC), FEV expression shows a similar alteration pattern to AR activity during the early stages of disease progression (Figure 6D). Consistently, in LNCaP xenografts, castration resulted in a significant reduction in AR activity accompanied by decreased FEV expression (Figure 6E). Experimental modulation of AR signaling in 22Rv1 cells further supports this association. Dihydrotestosterone (DHT) treatment, which enhanced AR activity, increased FEV expression, whereas knockdown of either full-length AR (AR-FL) or the AR splice variant AR-V7 resulted in reduced FEV expression (Figure 6F). Conversely, FEV knockdown in 22Rv1 cells is also accompanied by decreased expression of AR and canonical AR target genes (Figure 6G). Collectively, these findings suggest a potential reciprocal relationship between AR signaling and FEV expression during primary prostate cancer progression, and indicate that FEV may contribute to the maintenance of AR-associated transcriptional programs in castration-resistant prostate cancer.

Moreover, the relationship between FEV expression and response to AR-targeted therapy was examined. Although AR activity did not differ significantly between enzalutamide-sensitive and enzalutamide-resistant samples, FEV expression is significantly higher in the enzalutamide-sensitive group (Figure 6E). Consistently, in cell line models treated with enzalutamide or abiraterone, FEV expression is lower in treatment-resistant cells (Figure S7B). Thus, FEV expression may provide information beyond AR activity itself and potentially serve as a biomarker for therapeutic response. Consistent with its potential clinical relevance, lower FEV expression is associated with poorer survival outcomes in prostate cancer patients (Figure 6H).

## 3. Discussion

This study identifies a MIC-like tumor cell state associated with metastatic potential and highlights FEV as a candidate regulator associated with this phenotype. FEV is associated with a hybrid epithelial–mesenchymal state, and its perturbation disrupts this phenotype, particularly affecting epithelial and mesenchymal characteristics and reducing prostate cancer cell migration and invasion. In parallel, multi-omics and experimental evidence supports the regulation of MIF-family genes by FEV, while single-cell and spatial analyses link FEV-associated changes to reduced cytotoxicity and interferon responses in the tumor microenvironment. Together, these findings suggest that FEV may connect tumor cell state plasticity with microenvironmental remodeling during prostate cancer progression. Notably, FEV shows a reciprocal association with androgen signaling during primary prostate cancer progression and distinct expression patterns across treatment response groups, highlighting its potential relevance as a biomarker for disease progression and therapeutic response.

The existing literature on FEV in prostate cancer is limited, with one study suggesting a potential tumor-suppressive role for FEV in prostate cancer [13]. This study reported that FEV expression was reduced in recurrent and metastatic prostate cancer samples and was negatively associated with Gleason scores and pathological stages. The present study is consistent with these observations and further extends the potential functions of FEV to the early stages of prostate cancer progression. Previous work showed that FEV overexpression in LNCaP and PC-3 cells reduces their migratory and invasive capacities. In contrast, the present study demonstrates that FEV may exert broader functions beyond a conventional tumor-suppressive role. Specifically, FEV depletion in 22Rv1 cells disrupts the maintenance of the MIC-like state and impairs migratory and invasive capacities. Together, the multi-omics analyses, single-cell resolution, and cell line experiments in this study further expand the understanding of the multifaceted role of FEV in prostate cancer progression.

Moreover, the positive association between FEV expression and AR activity during the early stages of primary prostate cancer suggests a potential context-dependent relationship between FEV and AR signaling. Notably, this association appears to become uncoupled during advanced and metastatic disease. In the scRNA-seq dataset, FEV expression is undetectable in tumor cells from all bone metastatic lesions, indicating a marked loss of FEV expression in the bone metastatic setting. Consistently, FEV expression is nearly undetectable in the bone-derived metastatic prostate cancer cell lines PC3 and MDA-PCa-2b and the brain-derived metastatic cell line DU145, irrespective of AR activity (Figure S8A). Similarly, FEV expression is markedly reduced in metastatic castration-resistant prostate cancer spatial transcriptomic specimens despite relatively high AR activity (Figure S8B). In contrast, the neuroendocrine prostate cancer cell line NCI-H660 exhibits relatively high FEV expression despite low AR activity (Figure S8A). Together, these observations indicate that the positive association between AR activity and FEV expression is context dependent and is more evident during primary tumor progression, whereas FEV expression may become uncoupled from AR signaling during metastatic progression and lineage divergence. These findings further suggest that the relationship between FEV and AR signaling may vary across disease stages and metastatic or lineage contexts. Further studies are warranted to determine the functional role of FEV in specific metastatic sites and disease states.

## 4. Material and Methods

### 4.1 Datasets involved in this study

Multiple publicly available transcriptomic and epigenomic datasets were analyzed to characterize the association of FEV with primary prostate cancer progression. The principal scRNA-seq dataset was obtained from Synapse (Synapse ID: syn62736616) [20], with additional scRNA-seq data obtained from a previously published study [21]. Bulk transcriptomic datasets included The Cancer Genome Atlas prostate adenocarcinoma cohort (TCGA-PRAD), the Genotype-Tissue Expression (GTEx) project, GSE6956, and GSE134051. Spatial transcriptomic data included GSE278936 and spatial transcriptomic datasets from Synapse (Synapse ID: syn62767130, syn62767174) [20], which generated using the 10x Genomics Visium platform (Visium_FFPE_Human_Prostate_Cancer and Visium_FFPE_Human_Prostate_IF). Transcriptomic data from LNCaP xenograft models were obtained from GSE211856. ATAC-seq data for 22Rv1 cells were obtained from SRR6933113 and SRR6933114, and publicly available ChIP-seq data were obtained from GSE74315. Prostate cancer cell-line expression profiles were obtained from CCLE. Additional treatment-associated prostate cancer cell-line datasets were obtained from GSE86503, GSE94013, GSE150807, GSE120005, and GSE78201.

### 4.2 Cell and subtype annotation for scRNA-seq data

The primary scRNA-seq dataset was annotated based on canonical cell-type markers (Figure S1A). To further distinguish malignant epithelial cells from non-malignant epithelial populations, CNV profiles were inferred using inferCNV [22], with CD4+ T cells, CD8+ T cells, B cells, and NK cells serving as reference populations. A cell-level CNV score was calculated as the mean squared deviation of the inferred CNV profile from the neutral state, represented by a value of 1. Following identification of malignant epithelial cells based on their CNV profiles (Figure S1C), primary prostate tumor cells were further clustered using Seurat with a resolution parameter of 0.8. The resulting tumor cell clusters were defined as tumor subtypes for downstream analyses.

### 4.3 Functional cell state annotation

Following cell and tumor-subtype annotation, tumor cells were further characterized in terms of their developmental trajectories, developmental potential, cell-cycle states, and functional activities. Tumor-cell developmental trajectories were inferred using Monocle [23]. Cellular developmental potential was evaluated using CytoTRACE2 [24], with higher scores indicating greater inferred developmental potential and cellular plasticity. Cell-cycle activity was assessed using the CellCycleScoring function in Seurat, and each cell was assigned to the G1, S, or G2/M phase based on its relative S- and G2/M-phase scores.

Functional states were characterized using gene set activity scores. For scRNA-seq and spatial transcriptomic data, gene set scores were calculated using AddModuleScore, whereas gene set variation analysis (GSVA) [25] was applied to bulk transcriptomic data. The gene sets used in this study were obtained from the Molecular Signatures Database (MSigDB, version 2025.1) [26], supplemented with custom gene sets for specific biological processes. The custom antigen-presentation gene set comprised HLA-A, HLA-B, HLA-C, B2M, TAP1, and TAP2 [27]. Cytotoxicity was assessed using a predefined gene set [28]. comprising KLRF1, GNLY, CTSW, NKG7, KLRD1, GZMA, ADGRG1, CST7, KLRK1, FASLG, HCST, KLRB1, ITGB1, GZMB, and PRF1. Androgen receptor (AR) activity was assessed using an AR target gene set [29] comprising KLK3, KLK2, FKBP5, STEAP1, STEAP2, PPAP2A, RAB3B, ACSL3, and NKX3-1. These functional scores were subsequently used to characterize the biological states of tumor-cell subtypes and their associations with disease progression.

To further characterize the molecular features of tumor-cell subtypes, differentially expressed genes were identified between specific tumor-cell clusters using FindMarkers, followed by functional enrichment analysis using WebGestalt [30]. Transcription factor regulatory programs were inferred using SCENIC [31]. For each regulon, regulon activity in each tumor-cell cluster was compared with that in all other tumor-cell subtypes using the Wilcoxon rank-sum test to identify cluster-enriched transcriptional regulatory programs.

Cell–cell communication within the tumor microenvironment was analyzed using CellChat [32]. Outgoing and incoming signaling strengths were quantified for each tumor cell subtype by summing the inferred communication probabilities. Ligand–receptor interactions involving specific tumor cell subtypes were ranked according to their inferred interaction probabilities. Pathway-level signaling was characterized by examining ligand–receptor interactions associated with specific signaling pathways. For each tumor cell subtype, the corresponding interaction probabilities were summed to obtain an overall pathway-specific signaling score.

### 4.4 Mapping metastatic tumor cells to primary tumor subtypes

To characterize the transcriptional similarity between metastatic tumor cells and primary tumor cell subtypes, primary and metastatic epithelial tumor cells were jointly normalized, followed by variable feature selection, scaling, and principal component analysis (PCA). The first 30 principal components were used to represent the transcriptional state of individual cells. For each metastatic tumor cell, the nearest primary tumor cell in the PCA space was identified using a k-nearest neighbors (KNN) approach. The tumor-cell subtype of the matched primary tumor cell was then assigned to the corresponding metastatic tumor cell as its most transcriptionally similar primary tumor subtype. The proportions of metastatic tumor cells assigned to each primary tumor subtype were subsequently calculated to characterize the distribution of metastatic cells across primary tumor-cell states.

### 4.5 Spatial transcriptomic analysis

For spatial transcriptomic data, epithelial tumor spots were identified based on a positive epithelial gene set score calculated from EPCAM, KRT8, KRT18, and KRT19. The spatial distribution and abundance of specific immune cell populations were estimated using Cell2location [33]. Spatial neighborhood relationships between spots were determined using KNN method based on the spatial coordinates of each spot. Tumor boundary spots were defined as epithelial-enriched spots that had at least one non-epithelial spot among their six nearest spatial neighbors. For each epithelial spot, the Euclidean distance to the nearest tumor boundary spot was subsequently calculated to quantify its relative spatial position within the tumor region. These spatial features were used to characterize the localization of tumor-cell states and their spatial associations with the surrounding immune microenvironment.

### 4.6 Transcription factor regulatory relationship analysis

Potential TF regulatory relationships at specific genomic loci were investigated by integrating ATAC-seq derived chromatin accessibility profiles, TF motif prediction, and publicly available ChIP-seq data. ATAC-seq datasets were downloaded from the Sequence Read Archive (SRA) using the SRA-Toolkit. Paired-end sequencing reads were converted to FASTQ format and aligned to the human GRCh37/hg19 reference genome using Bowtie2. The resulting BAM files were generated and sorted using Samtools. Read groups were assigned using Picard, followed by removal of PCR duplicates using Picard MarkDuplicates. Accessible chromatin regions were identified by peak calling with MACS2 [34] using the paired-end BAM files. TF binding motifs within accessible chromatin regions were identified using motif annotations from the JASPAR database. ChIP-seq peak data were subsequently integrated with the genomic annotation of the mouse gene locus to assess the concordance between predicted TF binding sites, chromatin accessibility, and experimentally supported TF occupancy.

### 4.7 Statistical analysis

Unless otherwise specified, comparisons between two independent groups were performed using the two-sided Wilcoxon rank-sum test. Associations between continuous variables were assessed using Spearman correlation test. Progression-free survival (PFS) was analyzed using the Kaplan–Meier method, and survival distributions were compared using the log-rank test. All statistical tests were two-sided, and *P* < 0.05 was considered statistically significant.

### 4.8 Experimental validation

The human prostate cancer cell lines 22Rv1 were bought from Bestway Biotechnology (Xi’ an, China). Small interfering RNAs (siRNAs) targeting FEV were designed against the human FEV mRNA sequence. Three independent duplexes directed against non-overlapping regions of the transcript were used in parallel to control for sequence-specific off-target effects:

siRNA-FEV-135: CCCUGCUGAUCAACAUGUATT

siRNA-FEV-422: CUGCGCUACUACUACGACATT

siRNA-FEV-992: CUUGUCUGGUACUAGAUUUTT.

A non-targeting scrambled siRNA (siNC) served as the negative control and a siRNA targeting GAPDH as the positive transfection control. All siRNA were synthesized by GenePharma (Shanghai, China). Transient transfection was performed with Lipofectamine™ 3000 Transfection Reagent (Thermo Fisher Scientific; cat. no. L3000001) according to the manufacturer’s instructions, with minor modifications. For each well of a 6-well plate, 20 pmol of siRNA (final concentration, 20 nM) was diluted in 125 µL of Opti-MEM™ I Reduced-Serum Medium (Gibco), and, in a separate tube, 5 µL of Lipofectamine™ 3000 was diluted in 125 µL of Opti-MEM™. The two solutions were combined (1:1), mixed gently by pipetting, and incubated for 10 – 15 min at room temperature (20–25 °C) to allow the formation of siRNA–lipid complexes. The complexes (250 µL per well) were then added dropwise onto the cells, which had been fed with 1.75 mL of fresh, pre-warmed, antibiotic-free complete medium, and the plate was rocked gently to ensure even distribution. Cells were incubated at 37 °C with 5% CO2 for 6 h, after which the transfection medium was replaced with complete medium containing antibiotics. Cells were cultured for a further 42–66 h and harvested at 72 h (protein analysis) post-transfection.

Cells were lysed in RIPA lysis buffer (NCMBiotech, WB3100) supplemented with protease and phosphatase inhibitors (NCMBiotech, P002) and incubated on ice for 15 min. Lysates were centrifuged at 12,000 rpm for 15 min at 4 ° C, and the protein-containing supernatants were collected. Equal amounts of protein were mixed with SDS–PAGE sample loading buffer (Beyotime, P0015) and denatured by boiling at 100 °C for 10 min. Proteins were separated on YoungPAGE gels (GenScript, M00930) using MOPS running buffer and subsequently transferred onto PVDF membranes (Millipore, IPVH00010) using a wet transfer system. Membranes were blocked with PVA to prevent non-specific binding and incubated overnight at 4 °C with primary antibodies. After washing, membranes were incubated with the corresponding secondary antibodies for 2 h at room temperature. Protein bands were visualized using the e-BLOT imaging system. Primary antibodies used included anti-human FEV (Proteintech, 13437-1-AP, 1:2000), anti-human AR (Proteintech, 81844-1-RR, 1:2000), anti-human KLK3 (Proteintech, 84059-1-RR, 1:2000), anti-human TMPRSS2 (Proteintech, 80152-1-RR, 1:2000), anti-human NKX3.1 (Proteintech, 86085-1-RR, 1:2000), anti-human MIF (Proteintech, 83199-1-RR, 1:2000), anti-human DDT (Proteintech, 12389-1-AP, 1:2000)and anti- β -tubulin (Proteintech, HRP-60008, 1:5000). The secondary antibody used was anti-rabbit IgG (Cell Signaling Technology, 7074S, 1:5000).

Wound healing and Transwell assays were performed to assess cell migratory capacity. For the Transwell migration assay, cells were seeded into 24-well Transwell chambers (Corning, NY, USA) at a density of 5 × 10⁴ cells per well in 200 μL serum-free medium. After incubation, migrated cells were fixed with 4% paraformaldehyde solution and stained with 0.25% crystal violet. For the wound healing assay, transfected prostate cancer cells were cultured in six-well plates until approximately 80% density. A linear wound was created by vertically scratching using a 200 μL pipette, followed by replacement with 2 mL serum-free medium. Wound closure was monitored and quantified by measuring the average wound width over time.

## 5. Data availability

The single-cell transcriptomic datasets analyzed in this study include GSE193337, GSE143791, and the dataset associated with PMID: 34936871. Bulk transcriptomic datasets include The Cancer Genome Atlas (TCGA), Genotype-Tissue Expression (GTEx), GSE6956, and GSE134051. Spatial transcriptomic datasets include GSE278936 and two spatial transcriptomic datasets generated using the 10x Genomics platform (Visium_FFPE_Human_Prostate_Cancer and Visium_FFPE_Human_Prostate_IF). Cell line data were obtained from the Cancer Cell Line Encyclopedia (CCLE). Cell line datasets with treatment information were obtained from GSE86503, GSE94013, GSE150807, GSE120005, and GSE78201. LNCaP xenograft data were obtained from GSE211856. ATAC-seq data for 22Rv1 cells were obtained from SRR6933113 and SRR6933114, and ChIP-seq data were obtained from GSE74315.

## 6. Code availability

All code associated with this study is available at https://github.com/swu13/PRCAFEV.

## 7. Supplementary Data

Supplementary files are available online along with the manuscript.

## 8. Acknowledgement

This work was supported by the Fundamental Research Funds for the Central Universities and the National Natural Science Foundation of China (Grant No. 82227802, 62002270).

## Reference

[1] Raychaudhuri, R., Lin, D.W., Montgomery, R.B. (2025). Prostate Cancer: A Review. JAMA 333, 1433–1446.

[2] De Visser, K.E., Joyce, J.A. (2023). The evolving tumor microenvironment: From cancer initiation to metastatic outgrowth. Cancer Cell 41, 374–403.

[3] Bian, X., Wang, W., Abudurexiti, M., Zhang, X., Ma, W., Shi, G., Du, L., Xu, M., Wang, X., Tan, C., Sun, H., He, X., Zhang, C., Zhu, Y., Zhang, M., Ye, D., Wang, J. (2024). Integration Analysis of Single-Cell Multi-Omics Reveals Prostate Cancer Heterogeneity. Advanced science (Weinheim, Baden-Wurttemberg, Germany) 11, e2305724.

[4] De Vargas Roditi, L., Jacobs, A., Rueschoff, J.H., Bankhead, P., Chevrier, S., Jackson, H.W., Hermanns, T., Fankhauser, C.D., Poyet, C., Chun, F., Rupp, N.J., Tschaebunin, A., Bodenmiller, B., Wild, P.J. (2022). Single-cell proteomics defines the cellular heterogeneity of localized prostate cancer. Cell reports. Medicine 3, 100604.

[5] Luo, Y., Zhong, H., Shang, T., Hu, B., Yuan, D., Jia, X., Lin, R., Wang, Z., Fang, Y., Zhu, G., Song, J., Liu, Z., Yan, B., Sun, F., Jia, Z., Yu, Y., Mao, L., Huang, H., Zhu, J. (2026). Single-Cell and Spatial Transcriptomics Uncover Immune Dynamics and Cellular Heterogeneity in Benign Prostatic Hyperplasia and Prostate Cancer Transition. MedComm 7, e70760.

[6] Chukhu, M., Dahiya, U.R., Heemers, H.V. (2025). Evolving roles for the androgen receptor and its protein interactome in castration-resistant prostate cancer. Oncogene 44, 3883–3894.

[7] Han, H., Wang, Y., Curto, J., Gurrapu, S., Laudato, S., Rumandla, A., Chakraborty, G., Wang, X., Chen, H., Jiang, Y. (2022). Mesenchymal and stem-like prostate cancer linked to therapy-induced lineage plasticity and metastasis. Cell Rep. 39.

[8] Ge, R., Wang, Z., Cheng, L. (2022). Tumor microenvironment heterogeneity an important mediator of prostate cancer progression and therapeutic resistance. NPJ precision oncology 6, 31.

[9] Keshavarzian, T., Furlano, K., Grillo, G., Mout, L., Arlidge, C., Hasan, F., Nand, A., Mikutenaite, M., Karadoulama, E., Goyal, A. (2025). Prostate cancer cells converge to an inflammatory-like state upon metastatic dissemination. Nat Commun. 16, 11339.

[10] Jin, H.-J., Zhao, J.C., Wu, L., Kim, J., Yu, J. (2014). Cooperativity and equilibrium with FOXA1 define the androgen receptor transcriptional program. Nat Commun. 5, 3972.

[11] Chen, Z., Wu, D., Thomas-Ahner, J.M., Lu, C., Zhao, P., Zhang, Q., Geraghty, C., Yan, P.S., Hankey, W., Sunkel, B. (2018). Diverse AR-V7 cistromes in castration-resistant prostate cancer are governed by HoxB13. Proceedings of the National Academy of Sciences 115, 6810–6815.

[12] Zhou, T., Yu, C., Han, Y., He, B., Feng, Q. (2025). GATA2 up-regulation restores androgen receptor chromatin association and advances darolutamide resistance in prostate cancer. Genes & Diseases 12, 101508.

[13] Liang, Y.X., Liang, Y.K., Zou, Z.H., Zhuo, Y.J., Ye, J.H., Zhu, X.J., Cai, Z.D., Lin, Z.Y., Mo, R.J., Wu, S.L., Zhang, Y.Q., Zhong, W.D. (2022). Tumor Suppressor Role and Clinical Significance of the FEV Gene in Prostate Cancer. Dis. Markers 2022, 8724035.

[14] Massagué, J., Ganesh, K. (2021). Metastasis-Initiating Cells and Ecosystems. Cancer Discov. 11, 971–994.

[15] Fares, J., Fares, M.Y., Khachfe, H.H., Salhab, H.A., Fares, Y. (2020). Molecular principles of metastasis: a hallmark of cancer revisited. Signal Transduct Target Ther 5, 28.

[16] Ledesma-Bazan, S., Sanchis, P., Seniuk, R., Pascual, G., Rada, M.J., Agulleiro, M., Russo, C., Valacco, P., Vazquez, E., Gueron, G. (2026). PCGEM1 overload: Triggering lethal stress responses in castration-resistant prostate cancer. Cancer Res. 86, 5908–5908.

[17] Merk, M., Zierow, S., Leng, L., Das, R., Du, X., Schulte, W., Fan, J., Lue, H., Chen, Y., Xiong, H. (2011). The D-dopachrome tautomerase (DDT) gene product is a cytokine and functional homolog of macrophage migration inhibitory factor (MIF). Proceedings of the National Academy of Sciences 108, E577–E585.

[18] Ovek Baydar, D., Rauluseviciute, I., Aronsen, D.R., Blanc-Mathieu, R., Bonthuis, I., De Beukelaer, H., Ferenc, K., Jegou, A., Kumar, V., Lemma, R.B. (2026). JASPAR 2026: expansion of transcription factor binding profiles and integration of deep learning models. Nucleic Acids Res. 54, D184–D193.

[19] Jones, M.G., Sun, D., Min, K.H.J., Colgan, W.N., Wang, H., Török, T., Ribeiro, J., Xue, J., Cardoso, E.C., Rong, Y. (2026). Spatiotemporal lineage tracing reveals the dynamic spatial architecture of tumor growth and metastasis. Nat. Genet., 1-13.

[20] Wu, S., Zhang, J., Wang, Y., Qin, X., Zhang, Z., Lu, Z., Kim, P., Zhou, X., Huang, L. (2024). metsDB: a knowledgebase of cancer metastasis at bulk, single-cell and spatial levels. Nucleic Acids Res.

[21] Tuong, Z.K., Loudon, K.W., Berry, B., Richoz, N., Jones, J., Tan, X., Nguyen, Q., George, A., Hori, S., Field, S. (2021). Resolving the immune landscape of human prostate at a single-cell level in health and cancer. Cell Rep. 37.

[22] Tickle Timothy, Tirosh Itay, Georgescu Christophe, Brown Maxwell, Brian, H., 2019. inferCNV of the Trinity CTAT Project. Klarman Cell Observatory, Broad Institute of MIT and Harvard, Cambridge, MA, USA.

[23] Trapnell, C., Cacchiarelli, D., Grimsby, J., Pokharel, P., Li, S., Morse, M., Lennon, N.J., Livak, K.J., Mikkelsen, T.S., Rinn, J.L. (2014). The dynamics and regulators of cell fate decisions are revealed by pseudotemporal ordering of single cells. Nat. Biotechnol. 32, 381–386.

[24] Gulati, G.S., Sikandar, S.S., Wesche, D.J., Manjunath, A., Bharadwaj, A., Berger, M.J., Ilagan, F., Kuo, A.H., Hsieh, R.W., Cai, S., Zabala, M., Scheeren, F.A., Lobo, N.A., Qian, D., Yu, F.B., Dirbas, F.M., Clarke, M.F., Newman, A.M. (2020). Single-cell transcriptional diversity is a hallmark of developmental potential. Science 367, 405–411.

[25] Hänzelmann, S., Castelo, R., Guinney, J. (2013). GSVA: gene set variation analysis for microarray and RNA-seq data. BMC Bioinf. 14, 7.

[26] Liberzon, A., Birger, C., Thorvaldsdóttir, H., Ghandi, M., Mesirov, J.P., Tamayo, P. (2015). The Molecular Signatures Database (MSigDB) hallmark gene set collection. Cell Syst 1, 417–425.

[27] Rudin, C.M., Balli, D., Lai, W.V., Richards, A.L., Nguyen, E., Egger, J.V., Choudhury, N.J., Sen, T., Chow, A., Poirier, J.T., Geese, W.J., Hellmann, M.D., Forslund, A. (2023). Clinical Benefit From Immunotherapy in Patients With SCLC Is Associated With Tumor Capacity for Antigen Presentation. J. Thorac. Oncol. 18, 1222–1232.

[28] Wang, P.L., Lai, W.P., Zheng, J.M., Wu, X.F., Zhan, J.N., Yi, T.Z., Jin, Z.Y., Wu, X.L. (2025). Heterogeneous characteristics of γδ T cells in peripheral blood of diffuse large B-cell lymphoma. Biomarker research 13, 82.

[29] Spratt, D.E., Alshalalfa, M., Fishbane, N., Weiner, A.B., Mehra, R., Mahal, B.A., Lehrer, J., Liu, Y., Zhao, S.G., Speers, C., Morgan, T.M., Dicker, A.P., Freedland, S.J., Karnes, R.J., Weinmann, S., Davicioni, E., Ross, A.E., Den, R.B., Nguyen, P.L., Feng, F.Y., Lotan, T.L., Chinnaiyan, A.M., Schaeffer, E.M. (2019). Transcriptomic Heterogeneity of Androgen Receptor Activity Defines a de novo low AR-Active Subclass in Treatment Naïve Primary Prostate Cancer. Clin. Cancer Res. 25, 6721–6730.

[30] Elizarraras, J.M., Liao, Y., Shi, Z., Zhu, Q., Pico, A.R., Zhang, B. (2024). WebGestalt 2024: faster gene set analysis and new support for metabolomics and multi-omics. Nucleic Acids Res. 52, W415–W421.

[31] Aibar, S., González-Blas, C.B., Moerman, T., Huynh-Thu, V.A., Imrichova, H., Hulselmans, G., Rambow, F., Marine, J.C., Geurts, P., Aerts, J., van den Oord, J., Atak, Z.K., Wouters, J., Aerts, S. (2017). SCENIC: single-cell regulatory network inference and clustering. Nat. Methods 14, 1083–1086.

[32] Jin, S., Guerrero-Juarez, C.F., Zhang, L., Chang, I., Ramos, R., Kuan, C.H., Myung, P., Plikus, M.V., Nie, Q. (2021). Inference and analysis of cell-cell communication using CellChat. Nat Commun 12, 1088.

[33] Kleshchevnikov, V., Shmatko, A., Dann, E., Aivazidis, A., King, H.W., Li, T., Elmentaite, R., Lomakin, A., Kedlian, V., Gayoso, A., Jain, M.S., Park, J.S., Ramona, L., Tuck, E., Arutyunyan, A., Vento-Tormo, R., Gerstung, M., James, L., Stegle, O., Bayraktar, O.A. (2022). Cell2location maps fine-grained cell types in spatial transcriptomics. Nat. Biotechnol. 40, 661–671.

[34] Feng, J., Liu, T., Qin, B., Zhang, Y., Liu, X.S. (2012). Identifying ChIP-seq enrichment using MACS. Nat. Protoc. 7, 1728–1740.

