## Supplementary Figures1-8 for "An FEV-Associated Metastasis-Initiating-Like State Linked to Tumor Cell Plasticity and Immune Niche Remodeling in Prostate Cancer"

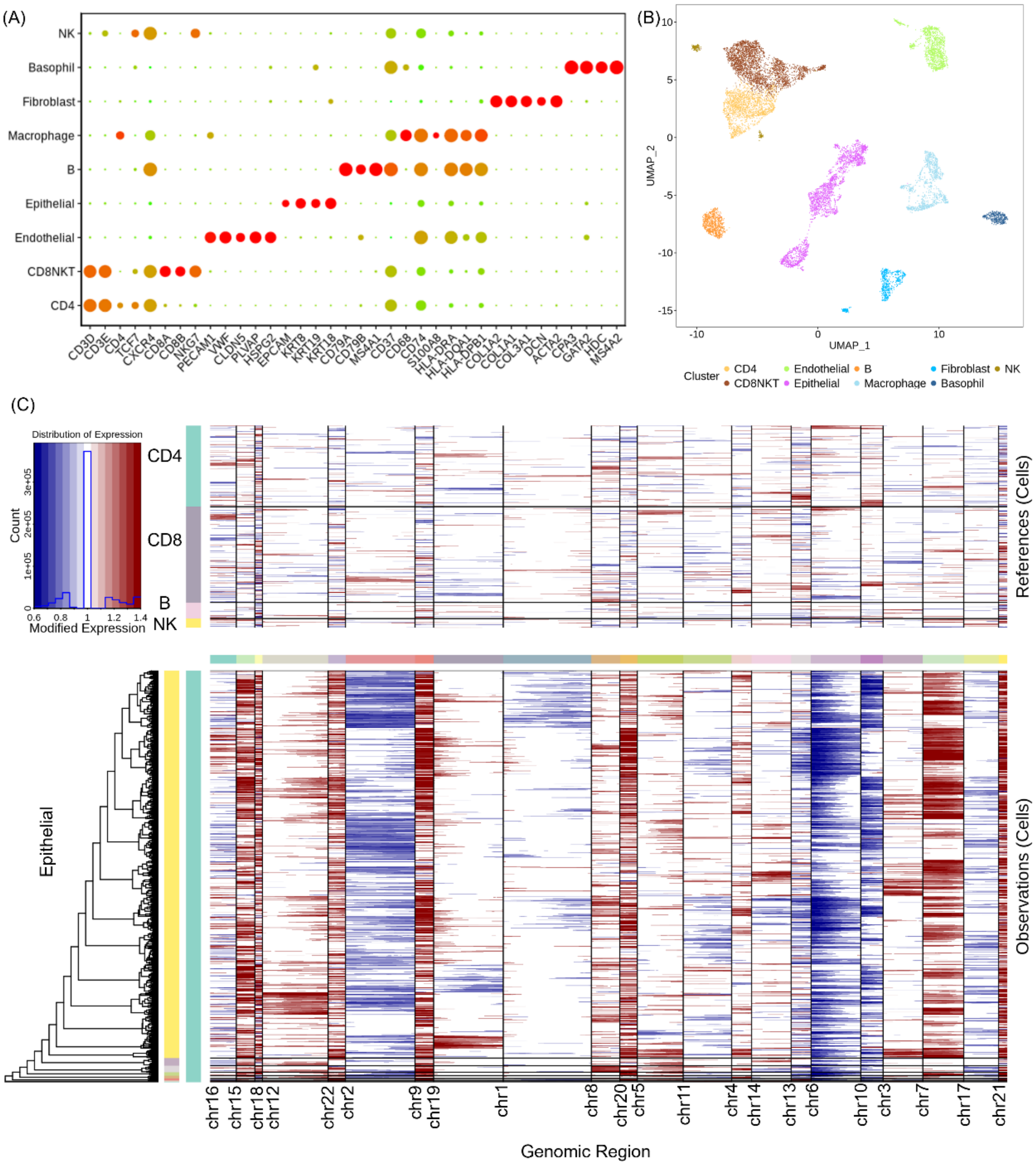


Figure S1. Identification of tumor cells in a prostate cancer single-cell dataset. (A) Cell type annotation based on canonical markers. (B) UMAP visualization of all the cell populations. (C) Inferred copy-number variation (CNV) profiles of epithelial cells identified in (A) using the inferCNV algorithm.

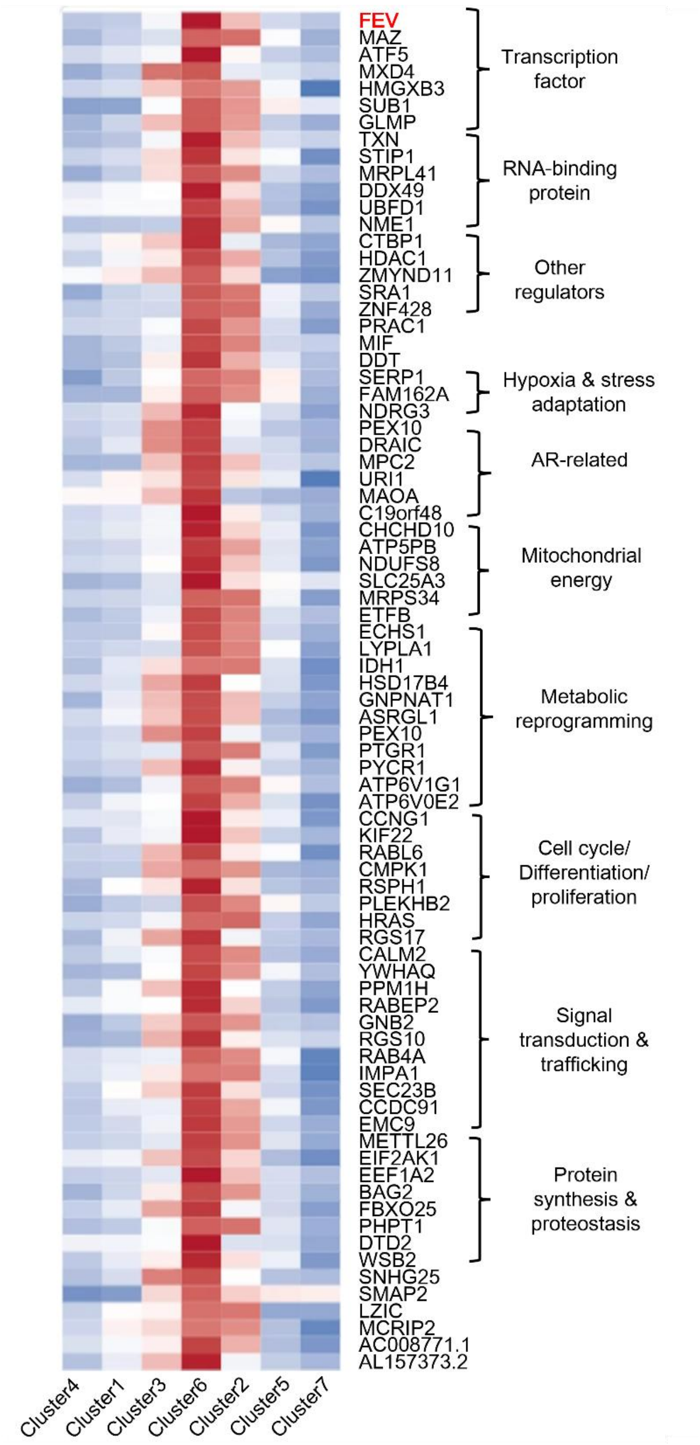


Figure S2. Trajectory-associated genes defining the cluster 6 tumor state. Genes showing increased expression from the trajectory root toward cluster 6 and subsequent decreases toward the terminal state.

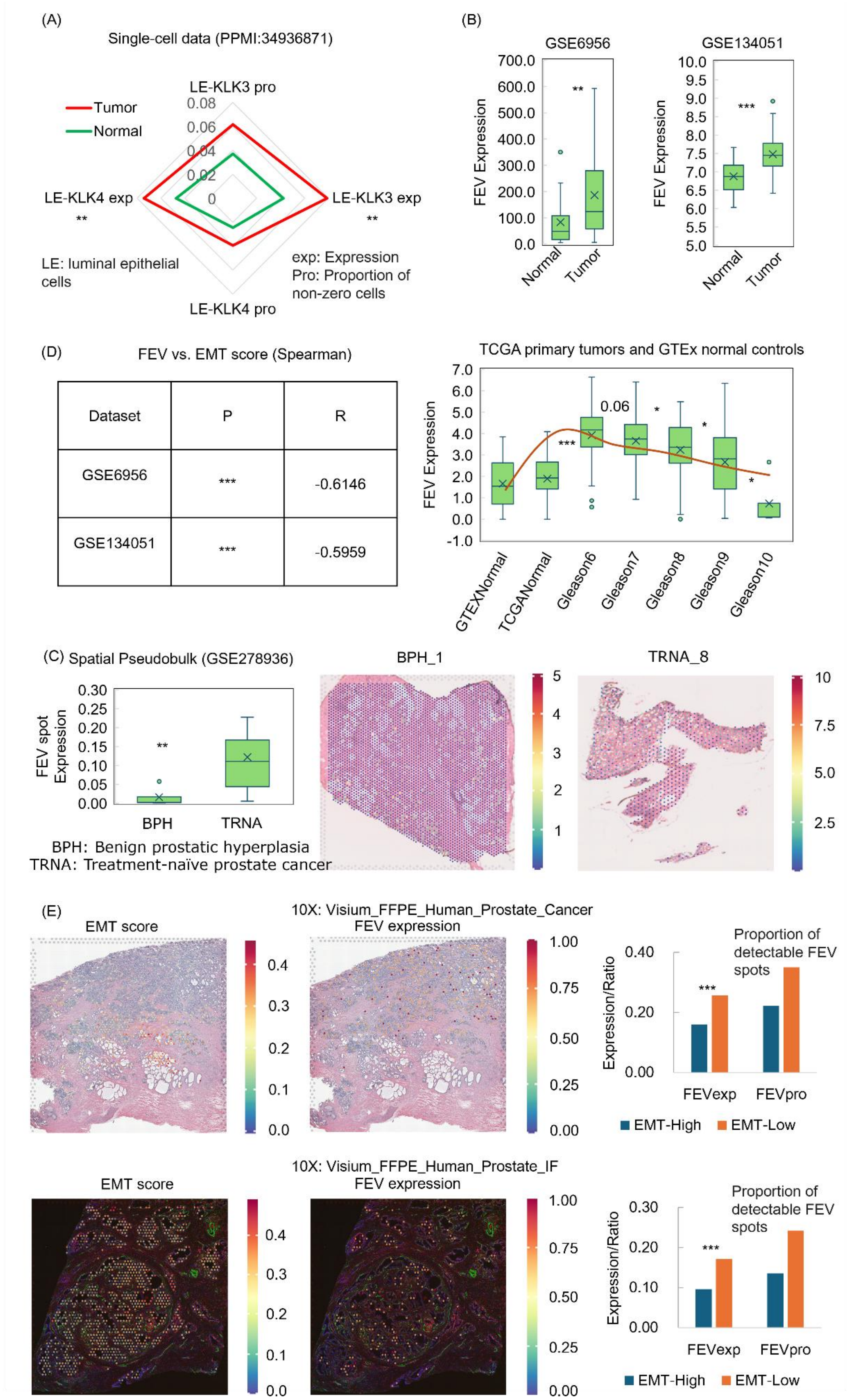


Figure S3. Association of FEV expression with prostate cancer malignancy and epithelial–mesenchymal transition (EMT). (A) Comparison of FEV expression and the proportion of cells with detectable FEV expression between tumor and normal luminal epithelial (LE) cells in a single-cell transcriptomic dataset. LE-KLK3 and LE-KLK4 denote LE cells with relatively high expression of KLK3 and KLK4, respectively. (B) Comparison of FEV expression between normal and tumor tissues in the GSE6956, GSE134051, The Cancer Genome Atlas (TCGA) and The Genotype-Tissue Expression (GTEx) datasets. (C) Comparison of pseudobulk FEV expression across spatial transcriptomic spots between benign prostatic hyperplasia (BPH) and treatment-naïve prostate cancer (TRAN), with representative spatial maps showing FEV expression in two tissue sections. (D) Spearman correlation between FEV expression and EMT scores in the GSE6956 and GSE134051 datasets. (E) Comparison of FEV expression and the proportion of spots with detectable FEV expression between spots with high and low EMT scores in two independent 10x Genomics spatial transcriptomic datasets.

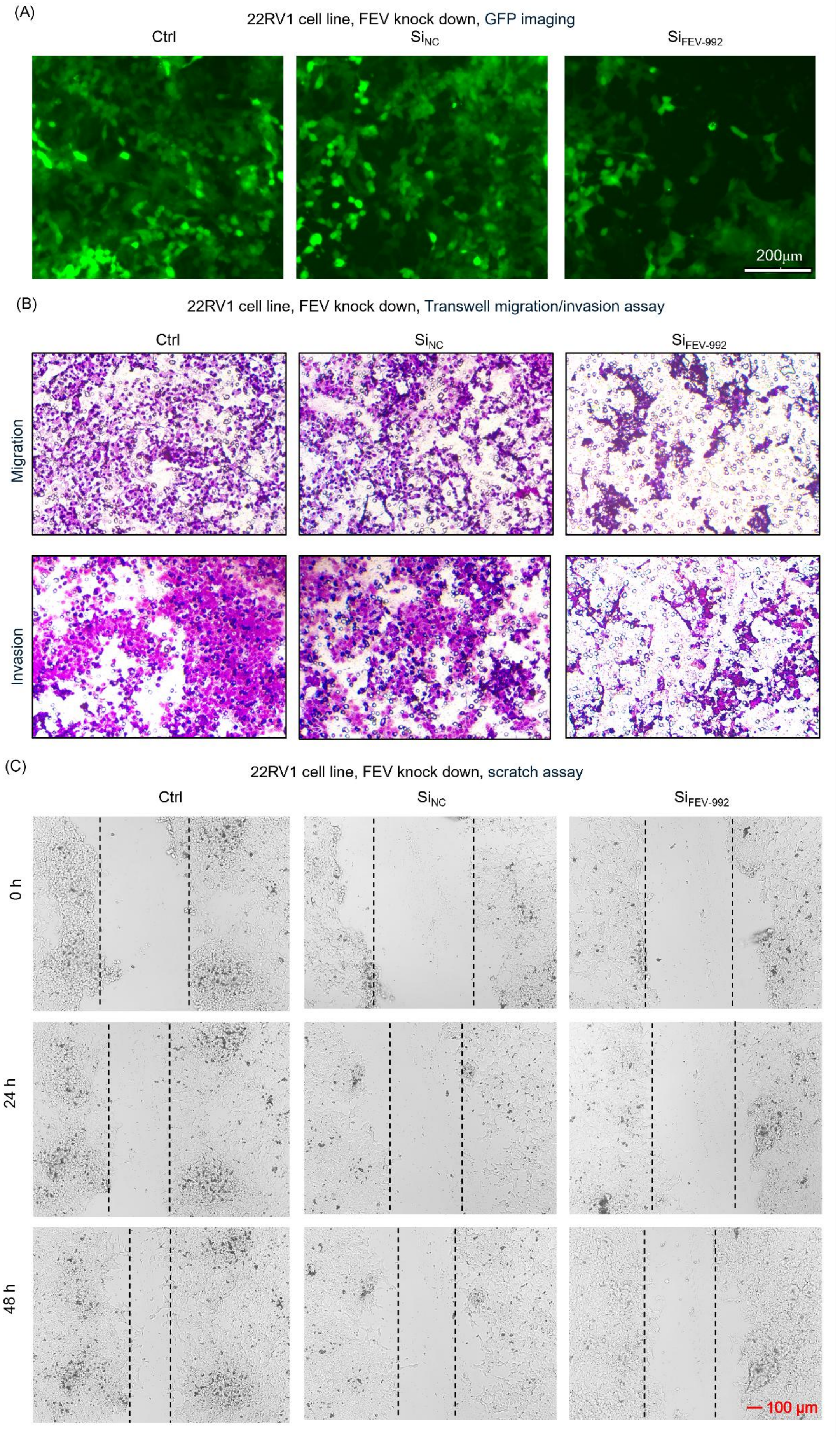

(A)
22RV1 cell line, FEV knock down, GFP imaging
Ctrl
$Si_{NC}$
$Si_{FEV-992}$
200μm
(B)
22RV1 cell line, FEV knock down, Transwell migration/invasion assay
Ctrl
$Si_{NC}$
$Si_{FEV-992}$
Migration
Invasion
(C)
22RV1 cell line, FEV knock down, scratch assay
Ctrl
$Si_{NC}$
$Si_{FEV-992}$
0 h
24 h
48 h
100 μm

Figure S4. Effects of FEV knockdown on the migratory and invasive capacities of 22Rv1 cells. (A) Representative GFP images of 22Rv1 cells following FEV knockdown. (B) Transwell migration and invasion assays of 22Rv1 cells following FEV knockdown. (C) Scratch-wound healing assays of 22Rv1 cells following FEV knockdown.

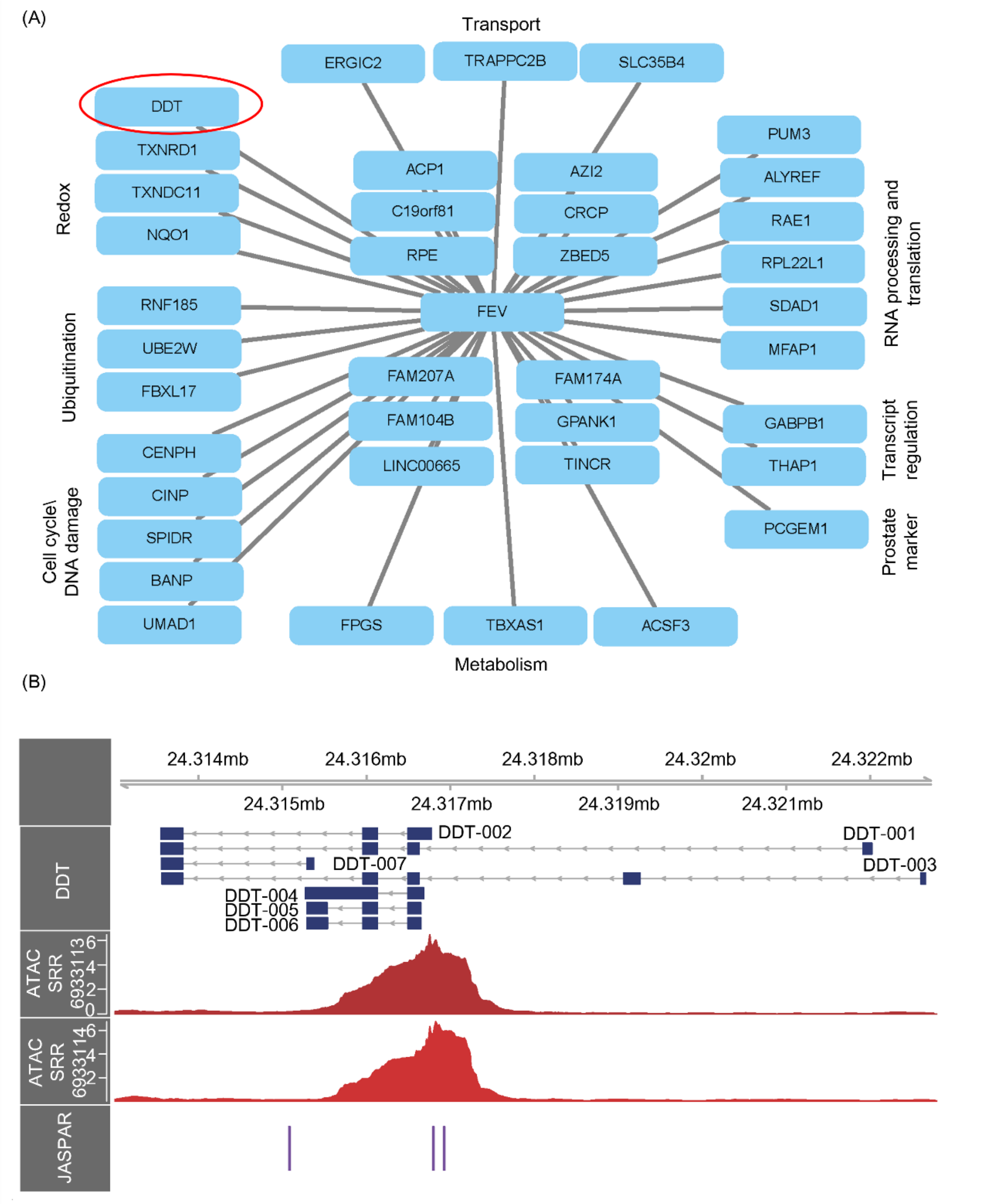


Figure S5. Downstream targets of FEV. (A) FEV target genes predicted by SCENIC analysis and their associated biological processes. (B) Accessible chromatin regions identified by ATAC-seq and putative FEV-binding sites predicted by JASPAR motif analysis at the DDT locus.

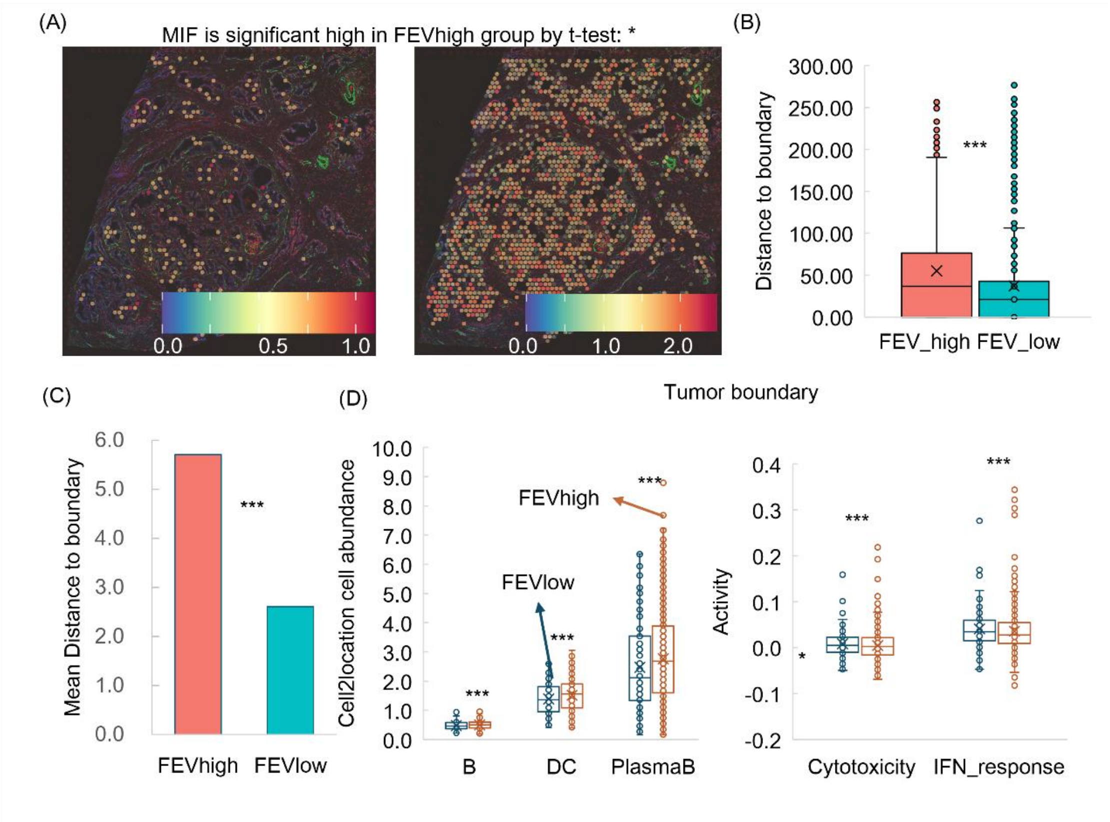


Figure S6. Associations of FEV expression with spatial immune remodeling. (A) Association between FEV and MIF expression within an independent spatial transcriptomic slide. (B-C) Comparison of the distance to the tumor boundary between FEV-high and FEV-low spots in two spatial transcriptomic slides. (D) Comparison of immune cell abundance and immune activity-associated scores between FEV-high and FEV-low spots at the tumor boundary within an independent spatial transcriptomic slide.

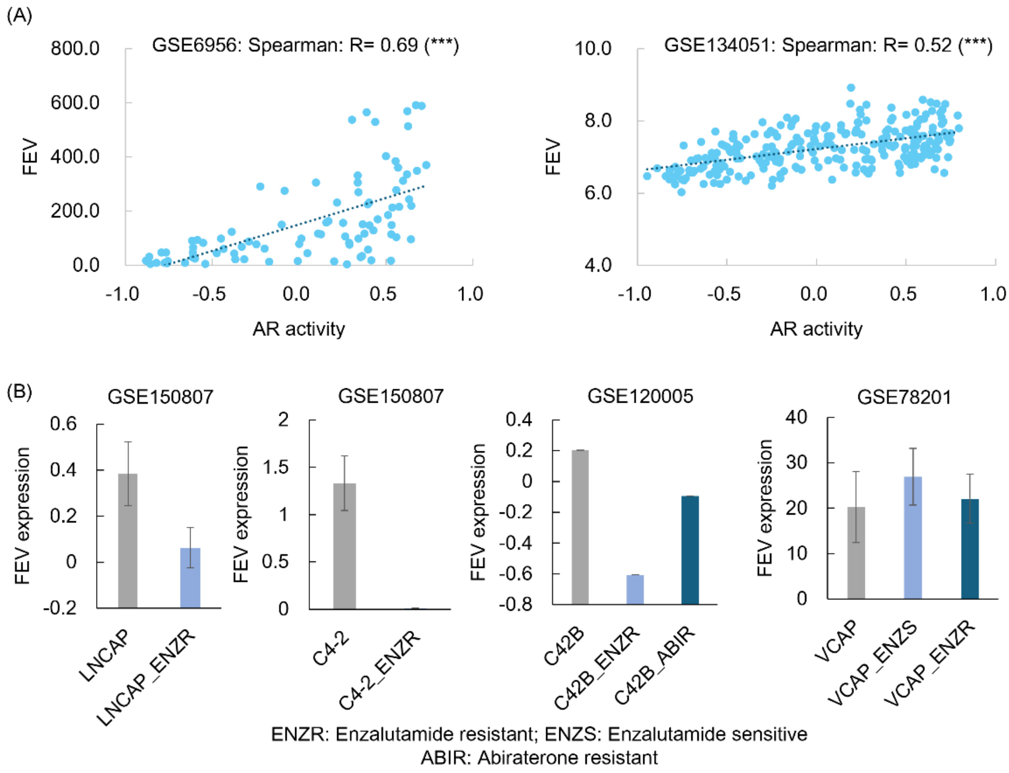


Figure S7. Associations between AR activity and FEV expression. (A) Associations between AR activity and FEV expression in two independent bulk transcriptomic datasets. (B) FEV expression of cell lines after drug treatment.

(A)

| Cell line | Location | AR | FEV |
|---|---|---|---|
| PC3 | Bone | -0.39 | 0 |
| MDAP CA2B | Bone | 0.59 | 0.03 |
| VCAP | Bone | 0.64 | 1.12 |
| DU145 | Brain | 0.23 | 0 |
| LNCA P | LN | 0.54 | 1.85 |
| NCIH6 60 | Neuroend ocrine prostate | -0.65 | 1.00 |

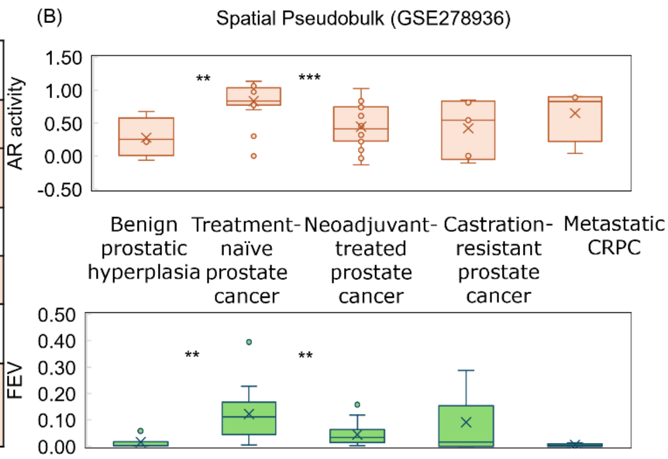


Figure S8. Context-dependent associations between FEV expression and AR activity. (A) FEV expression and AR activity across prostate cancer cell lines representing different disease and lineage contexts. (B) FEV expression and AR activity across different stages of prostate cancer in spatial transcriptomic datasets.
